# An acyltransferase protects *Caenorhabditis elegans* from thiol reductive stress through an autoinhibitory hypoxia response pathway

**DOI:** 10.1101/2025.10.09.681334

**Authors:** Ravi, Jogender Singh

**Affiliations:** Department of Biological Sciences, Indian Institute of Science Education and Research, Mohali, Punjab, 140306, India

**Keywords:** dithiothreitol, HIF-1, redox homeostasis, reductive stress, RHY-1

## Abstract

Cellular redox homeostasis depends on a finely tuned balance between oxidizing and reducing conditions, and disturbances in this balance lead to oxidative or reductive stress. While oxidative stress and its pathological outcomes are well studied, the molecular mechanisms underlying cellular responses to reductive stress remain poorly understood. Using *Caenorhabditis elegans* as a model, we investigate thiol reductive stress induced by dithiothreitol (DTT) and uncover a critical protective role for the hypoxia response pathway. We identify RHY-1, a membrane-associated acyltransferase and known negative regulator of the hypoxia-inducible factor HIF-1, as essential for survival under thiol reductive stress. Notably, *rhy-1* is a direct transcriptional target of HIF-1, and overexpression of *rhy-1* fully rescues the sensitivity of *hif-1* loss-of-function mutants to DTT. We demonstrate that RHY-1 functions in an autoinhibitory feedback loop, where elevated RHY-1 levels suppress activation of the hypoxia response pathway even during reductive stress. Together, our findings establish RHY-1 as both a regulatory and effector component of the hypoxia response pathway that mediates cellular protection against thiol reductive stress.

## Introduction

Redox homeostasis, the finely tuned balance between oxidizing and reducing environments, is essential for cellular survival (Ravi and Singh 2025). Reactive oxygen species (ROS), continuously generated as byproducts of metabolism, play crucial roles in cell signaling and homeostasis (Sies et al. 2022). However, excessive ROS accumulation leads to oxidative stress, which contributes to various pathological conditions, including neurodegeneration, cardiovascular disease, and cancer (Schieber and Chandel 2014; Forman and Zhang 2021). To counter oxidative insults, cells have evolved complex antioxidant defense systems comprising thioredoxin, glutathione (GSH), glutathione peroxidases, catalase, and superoxide dismutases (Kurutas 2016). In addition to endogenous systems, dietary antioxidants are frequently consumed under both physiological and pathological conditions to mitigate ROS-mediated damage (Kurutas 2016). Paradoxically, excessive production or intake of antioxidants can disrupt redox equilibrium, leading to reductive stress (Mercola 2025; Zhang et al. 2025). While oxidative stress responses have been extensively characterized, the mechanisms underlying reductive stress remain poorly understood (Ravi and Singh 2025).

Reductive stress has been implicated in several disorders, including cardiomyopathy, neurodegeneration, and metabolic dysfunction (Mercola 2025; Ravi and Singh 2025; Zhang et al. 2025). The limited understanding of reductive stress responses arises in part from the lack of robust experimental models capable of reliably inducing such conditions (Yang et al. 2022; Pan et al. 2024). Nevertheless, recent studies have begun to elucidate cellular mechanisms that sense and respond to reductive stress. For example, folliculin-interacting protein 1 (FNIP1) has been identified as a redox-sensitive molecule; its cysteine residues directly sense reductive stress, leading to proteasomal degradation of FNIP1 and subsequent increases in mitochondrial activity and ROS generation (Manford et al. 2020; Manford et al. 2021). In parallel, cells undergo extensive metabolic reprogramming under reductive stress to alleviate toxicity (Xiao and Loscalzo 2020; Yang et al. 2025).

Thiols, characterized by their sulfhydryl groups, constitute a major line of defense against oxidative stress by undergoing reversible oxidation to disulfides (Ulrich and Jakob 2019). However, excessive exposure to thiol-containing compounds, such as GSH, N-acetylcysteine, or dithiothreitol (DTT), can trigger reductive stress, adversely affecting development and lifespan in *Caenorhabditis elegans* (Gusarov et al. 2021; Gokul and Singh 2022; Winter et al. 2022; Ravi et al. 2023). DTT, traditionally thought to act primarily within the endoplasmic reticulum (ER) due to its oxidizing environment (Braakman et al. 1992; Lodish and Kong 1993; Tatu et al. 1993), has recently been shown to exert broader cellular effects. In addition to inducing ER stress, DTT can paradoxically generate oxidative stress by promoting futile redox cycling in the ER (Maity et al. 2016). Moreover, DTT perturbs the methionine-homocysteine cycle in *C. elegans* by upregulating the S-adenosylmethionine (SAM)-dependent methyltransferase RIPS-1 (Gokul and Singh 2022; Winter et al. 2022). This induction occurs via the hypoxia-inducible factor HIF-1-mediated hypoxia response pathway (Ravi et al. 2023). Although thiol reductive stress activates the hypoxia response, the physiological significance of this crosstalk remains unclear. It is not known whether activation of the hypoxia response represents a protective adaptation, nor have the downstream effector(s) mediating this protection been identified.

In this study, we demonstrated that the HIF-1-mediated hypoxia response pathway protects *C. elegans* from DTT-induced thiol stress. We further identified an acyltransferase, RHY-1, a transcriptional target of HIF-1, as a key protective factor against thiol stress. Interestingly, RHY-1 also functions as a negative regulator of the hypoxia pathway by inhibiting CYSL-1, a positive regulator of HIF-1 activation. Our data revealed that RHY-1 operates in an autoinhibitory feedback loop, where its overexpression suppresses hypoxia pathway activation even under thiol reductive stress conditions. Collectively, our findings establish RHY-1 as both a critical effector and regulator of the hypoxia response pathway during thiol reductive stress.

## Results

### The hypoxia response pathway protects *C. elegans* from thiol reductive stress

We previously reported that DTT upregulates the SAM-dependent methyltransferase RIPS-1 through the HIF-1-mediated hypoxia response pathway (Gokul and Singh 2022; Ravi et al. 2023). Elevated *rips-1* expression contributes to DTT toxicity, and *rips-1* loss-of-function mutants display resistance to 10 mM DTT (Gokul and Singh 2022). Since *hif-1* loss-of-function mutants lack induction of *rips-1* expression (Ravi et al. 2023), they were expected to exhibit similar resistance. However, *hif-1(ia4)* mutants were unable to develop on 10 mM DTT (Ravi et al. 2023). This observation suggested that HIF-1 may play a protective role under thiol reductive stress. Alternatively, the increased sensitivity of *hif-1* mutants could arise from residual basal *rips-1* expression, which is toxic in the presence of DTT.

To test whether basal *rips-1* expression accounts for the heightened DTT sensitivity of *hif-1(ia4)* mutants compared to *rips-1* mutants, we generated a *hif-1(ia4);rips-1(jsn11)* double mutant. The *rips-1(jsn11)* allele carries an early stop codon, resulting in a complete loss of functional RIPS-1 protein (Ravi et al. 2023). Therefore, *hif-1(ia4);rips-1(jsn11)* worms should lack any functional RIPS-1. As expected, *rips-1(jsn11)* single mutants developed normally on 10 mM DTT; however, *hif-1(ia4);rips-1(jsn11)* worms failed to develop (Fig. 1a-c). This result demonstrated that the increased sensitivity of *hif-1* mutants to DTT cannot be attributed to basal *rips-1* expression.

**Figure 1.**
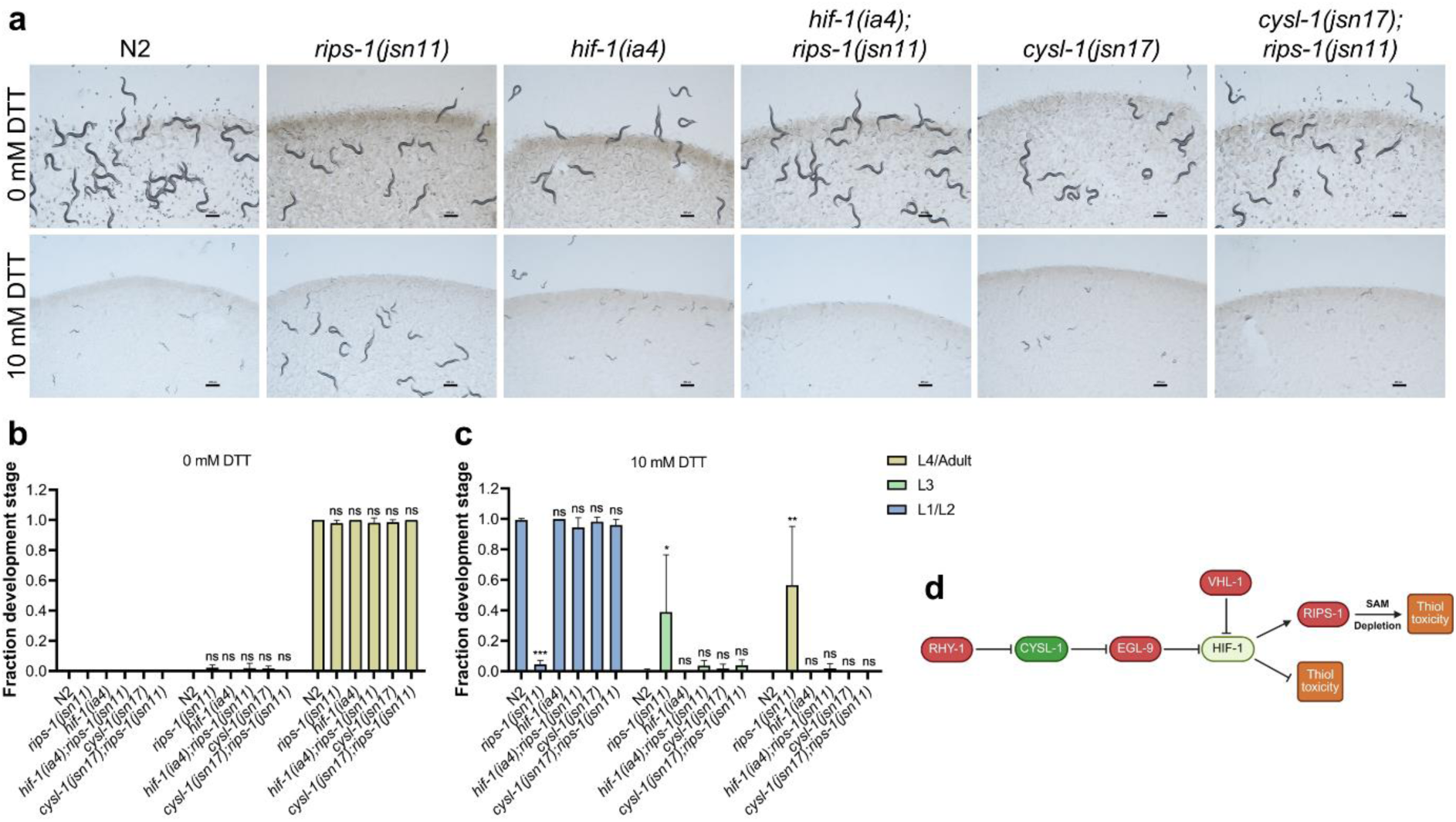
Hypoxia response pathway protects *C. elegans* from thiol reductive stress. (a) Representative images of N2, *rips-1(jsn11)*, *hif-1(ia4)*, *hif-1(ia4);rips-1(jsn11)*, *cysl-1(jsn17)*, and *cysl-1(jsn17);rips-1(jsn11)* worms on 0 mM and 10 mM DTT on an *E. coli* OP50 diet after 72 h of egg laying at 20°C. Scale bar = 200 μm. (b)-(c) Quantification of different developmental stages of N2, *rips-1(jsn11)*, *hif-1(ia4)*, *hif-1(ia4);rips-1(jsn11)*, *cysl-1(jsn17)*, and *cysl-1(jsn17);rips-1(jsn11)* worms on 0 mM (b) and 10 mM DTT (c) on an *E. coli* OP50 diet after 72 h of egg laying at 20°C. The different developmental stage symbols are shown in panel c. \*\*\**p* < 0.001, \*\**p* < 0.01, \**p* < 0.05, and non-significant (ns) via ordinary one-way ANOVA followed by Dunnett’s multiple comparisons test. For each developmental stage, the comparison is with N2 (*n* = 3 biological replicates; worms per condition per replicate > 70). (d) Schematic representation of the HIF-1-mediated hypoxia response pathway and its role in modulating thiol toxicity.

DTT is known to activate the hypoxia response pathway via the upstream regulator CYSL-1 (Fig. 1d). Similar to *hif-1(ia4)* worms, *cysl-1* loss-of-function mutants fail to induce *rips-1* expression and are sensitive to 10 mM DTT (Ravi et al. 2023). Consistently, *cysl-1(jsn17);rips-1(jsn11)* double mutants also failed to develop on 10 mM DTT (Fig. 1a-c). Together, these findings indicated that activation of the HIF-1-mediated hypoxia response pathway by DTT confers protection against thiol reductive stress (Fig. 1d).

### RHY-1 protects against thiol reductive stress

To further delineate the role of the hypoxia response pathway in defense against thiol reductive stress, we examined the effects of loss-of-function mutations in *rhy-1*, *egl-9*, and *vhl-1*. These genes are key regulators of the hypoxia response pathway and are required for the proteasomal degradation of the transcription factor HIF-1 (Epstein et al. 2001; Shen et al. 2006). Inactivation of any of these genes leads to stabilization of HIF-1 and constitutive activation of hypoxia-responsive genes (Shen et al. 2006). Because *rhy-1*, *egl-9*, and *vhl-1* mutants exhibit elevated *rips-1* expression (Ravi et al. 2023), they are expected to display increased sensitivity to DTT-induced reductive stress. To eliminate confounding effects arising from *rips-1* activity, we generated double mutants, *rhy-1(jsn13jsn14);rips-1(jsn11)*, *egl-9(sa307);rips-1(jsn11)*, and *vhl-1(jsn16);rips-1(jsn11)*, and assessed their sensitivity to 10 mM DTT. The alleles *rhy-1(jsn13jsn14)*, *egl-9(sa307)*, and *vhl-1(jsn16)* are loss-of-function mutations consisting of two missense substitutions (S146Y and R183C), a 243-bp deletion, and a premature nonsense mutation, respectively (Darby et al. 1999; Ravi et al. 2023). We selected 10 mM DTT because higher concentrations can introduce secondary effects related to ER stress (Gokul and Singh 2022). Unexpectedly, while *rips-1(jsn11)* animals developed normally on 10 mM DTT, *rhy-1(jsn13jsn14);rips-1(jsn11)* animals failed to develop (Fig. 2a-c). In contrast, *egl-9(sa307);rips-1(jsn11)* and *vhl-1(jsn16);rips-1(jsn11)* animals showed improved development relative to *rhy-1(jsn13jsn14);rips-1(jsn11)*. This finding was surprising, as *rhy-1* mutants were anticipated to exhibit enhanced tolerance to DTT due to increased activation of the hypoxia response pathway.

**Figure 2.**
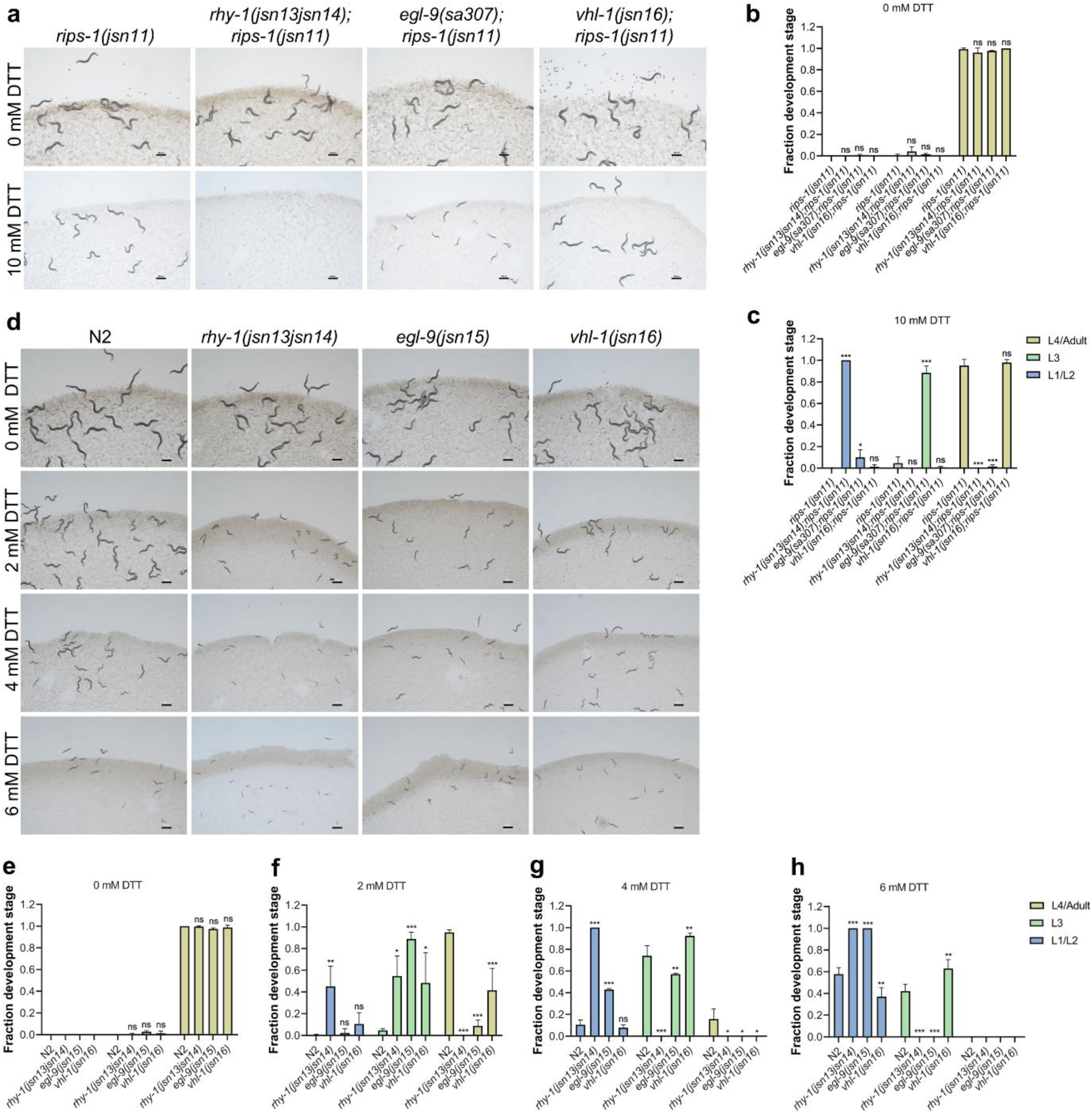
RHY-1 protects against thiol reductive stress. (a) Representative images of *rips-1(jsn11)*, *rhy-1(jsn13jsn14);rips-1(jsn11)*, *egl-9(sa307);rips-1(jsn11)*, and *vhl-1(jsn16);rips1(jsn11)* worms on 0 mM and 10 mM DTT on *E. coli* OP50 diet after 72 h of egg laying at 20°C. Scale bar = 200 μm. (b)-(c) Quantification of different developmental stages of *rips-1(jsn11)*, *rhy-1(jsn13jsn14);rips-1(jsn11)*, *egl-9(sa307);rips-1(jsn11)*, and *vhl-1(jsn16);rips1(jsn11)* worms on 0 mM (b) and 10 mM DTT (c) on *E. coli* OP50 diet after 72 h of egg laying at 20°C. The different developmental stage symbols are shown in panel c. \*\*\**p* < 0.001, \**p* < 0.05, and non-significant (ns) via ordinary one-way ANOVA followed by Dunnett’s multiple comparisons test. For each developmental stage, the comparison is with *rips-1(jsn11)* (*n* = 3 biological replicates; worms per condition per replicate > 75). (d) Representative images of N2, JSJ14 (*rhy-1(jsn13jsn14)*), JSJ15 (*egl-9(jsn15)*), and JSJ16 (*vhl-1(jsn16)*) worms on 0 mM, 2 mM, 4 mM, and 6 mM DTT on *E. coli* OP50 diet after 72 hours of egg laying. Scale bar = 200 μm. (e)-(h) Quantification of different developmental stages of N2, JSJ14 (*rhy-1(jsn13jsn14)*), JSJ15 (*egl-9(jsn15)*), and JSJ16 (*vhl-1(jsn16)*) worms on 0 mM (e), 2 mM (f), 4 mM (g), and 6 mM (h) DTT on *E. coli* OP50 diet after 72 hours of egg laying. The different developmental stage symbols are shown in panel h. \*\*\**p* < 0.001, \*\**p* < 0.01, \**p* < 0.05, and non-significant (ns) via ordinary one-way ANOVA followed by Dunnett’s multiple comparisons test. For each developmental stage, the comparison is with N2 (*n* = 3 biological replicates; worms per condition per replicate > 70).

To rule out nonspecific effects from the *rips-1(jsn11)* background, we next evaluated the DTT sensitivity of single mutants *rhy-1(jsn13jsn14)*, *egl-9(jsn15)*, and *vhl-1(jsn16)* across a range of DTT concentrations. The *egl-9(jsn15)* allele is a loss-of-function mutation caused by a splice acceptor site disruption (Ravi et al. 2023). As expected, all three mutants were more sensitive to DTT than wild-type N2 worms, consistent with elevated *rips-1* expression (Fig. 2d-h). However, *rhy-1(jsn13jsn14)* worms exhibited significantly greater sensitivity than either *egl-9(jsn15)* or *vhl-1(jsn16)* worms (Fig. 2d-h), suggesting a unique protective role for RHY-1 against DTT toxicity.

DTT toxicity can be alleviated by vitamin B12 supplementation, which restores SAM levels depleted by increased *rips-1* expression (Gokul and Singh 2022; Winter et al. 2022). We therefore tested whether vitamin B12 supplementation could rescue the DTT sensitivity of mutants with constitutively active hypoxia signaling. While vitamin B12 supplementation effectively rescued DTT-induced toxicity in N2, *egl-9(jsn15)*, and *vhl-1(jsn16)* worms, it failed to ameliorate the effects in *rhy-1(jsn13jsn14)* mutants (Supplementary Fig. 1a-c). Consistently, vitamin B12 also failed to rescue DTT toxicity in an independent *rhy-1* allele, *rhy-1(ok1402)* (Supplementary Fig. 1d-f). Taken together, these results demonstrated that RHY-1 protects from the toxic effects of DTT.

### RHY-1 functions downstream of HIF-1 to protect from thiol toxicity

The *rhy-1* gene encodes an acyltransferase that not only regulates HIF-1 activity but is itself a transcriptional target of HIF-1 (Shen et al. 2006). Consistent with this, *rhy-1* expression increases upon DTT exposure in a HIF-1-dependent manner (Ravi et al. 2023). Given that RHY-1 protects against DTT-induced toxicity, we investigated whether the protective role of HIF-1 under thiol reductive stress is mediated through RHY-1. To test this hypothesis, we carried out *rhy-1* rescue experiments. First, we examined whether overexpression of *rhy-1* under its native promoter (*rhy-1p::rhy-1*) could rescue the DTT sensitivity of *rhy-1(jsn13jsn14)* mutants. Indeed, *rhy-1* overexpression restored normal development of *rhy-1(jsn13jsn14)* worms on 10 mM DTT in the presence of 500 nM vitamin B12 (Supplementary Fig. 2a and b).

We next introduced the *rhy-1p::rhy-1* transgene into *hif-1(ia4)* mutants and assessed their development under thiol stress. *hif-1(ia4)* worms overexpressing *rhy-1* exhibited markedly improved development on 10 mM DTT compared to non-transgenic *hif-1(ia4)* worms (Fig. 3a and b). Similarly, supplementation with 500 nM vitamin B12 further enhanced their development relative to non-transgenic controls (Fig. 3a and b). Notably, *rhy-1* overexpression rescued development in *hif-1(ia4)* but not in *rhy-1(jsn13jsn14)* animals on 10 mM DTT (Fig. 3 and Supplementary Fig. 2). This difference is likely due to the HIF-1 dependence of *rips-1* expression, such that *hif-1(ia4)* animals may express lower levels of *rips-1* than *rhy-1(jsn13jsn14)* animals under *rhy-1* overexpression conditions. Together, these results demonstrated that *rhy-1* overexpression fully rescues the DTT sensitivity of *hif-1* mutants, indicating that HIF-1 confers protection against thiol reductive stress primarily through the induction of RHY-1.

**Figure 3.**
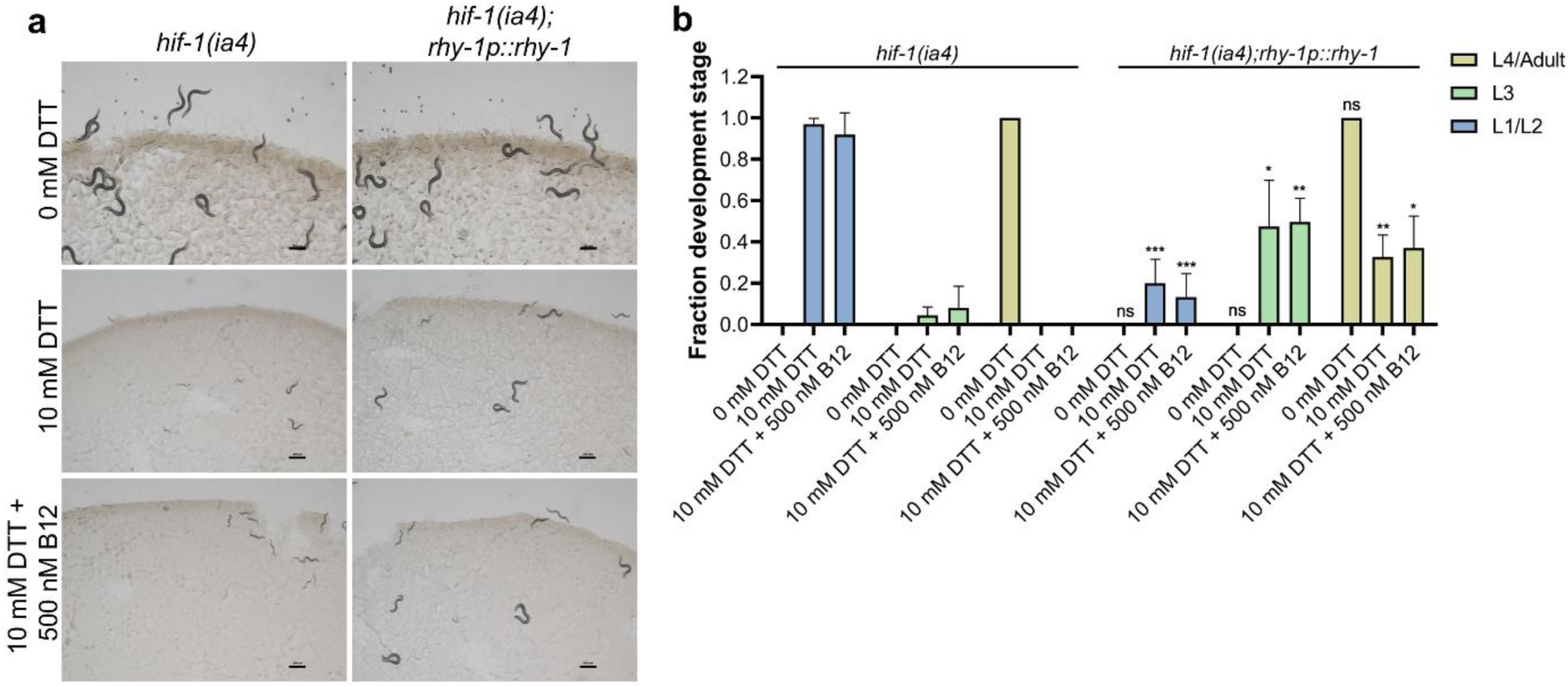
RHY-1 functions downstream of HIF-1 to protect from thiol toxicity. (a) Representative images of *hif-1(ia4)* and *hif-1(ia4);rhy-1p::rhy-1* worms exposed to 0 mM DTT, 10 mM DTT, or 10 mM DTT supplemented with 500 nM vitamin B12 on *E. coli* OP50 for 72 h at 20°C. Scale bar = 200 μm. (b) Quantification of different developmental stages of *hif-1(ia4)* and *hif-1(ia4);rhy-1p::rhy-1* worms exposed to 0 mM DTT, 10 mM DTT, or 10 mM DTT supplemented with 500 nM vitamin B12 on *E. coli* OP50 for 72 h at 20°C. \*\*\**p* < 0.001, \*\**p* < 0.01, \**p* < 0.05, and non-significant (ns) via the unpaired *t*-test. For each condition of *hif-1(ia4);rhy-1p::rhy-1* worms, the comparison is with the corresponding *hif-1(ia4)* condition (*n* = 3 biological replicates; worms per condition per replicate > 65).

### RHY-1 functions in an autoinhibitory feedback loop to protect from thiol toxicity

Previous studies have shown that *rhy-1* participates in a negative feedback loop that represses the targets of the hypoxia response pathway (Shen et al. 2006). However, the functional significance of *rhy-1* within this regulatory loop remained unclear. Our findings suggested that RHY-1 not only acts as a negative regulator of the hypoxia response pathway but also serves as an effector protein that mitigates thiol-induced toxicity. To test whether *rhy-1* overexpression can repress hypoxia pathway activation even under conditions that normally induce it, we overexpressed *rhy-1p::rhy-1* in *rips-1p::GFP* reporter worms. Adult worms were exposed to varying concentrations of DTT for 12 hours, followed by measurement of green fluorescent protein (GFP) levels. Strikingly, *rhy-1* overexpression completely suppressed the DTT-induced upregulation of *rips-1* (Fig. 4a and b).

**Figure 4.**
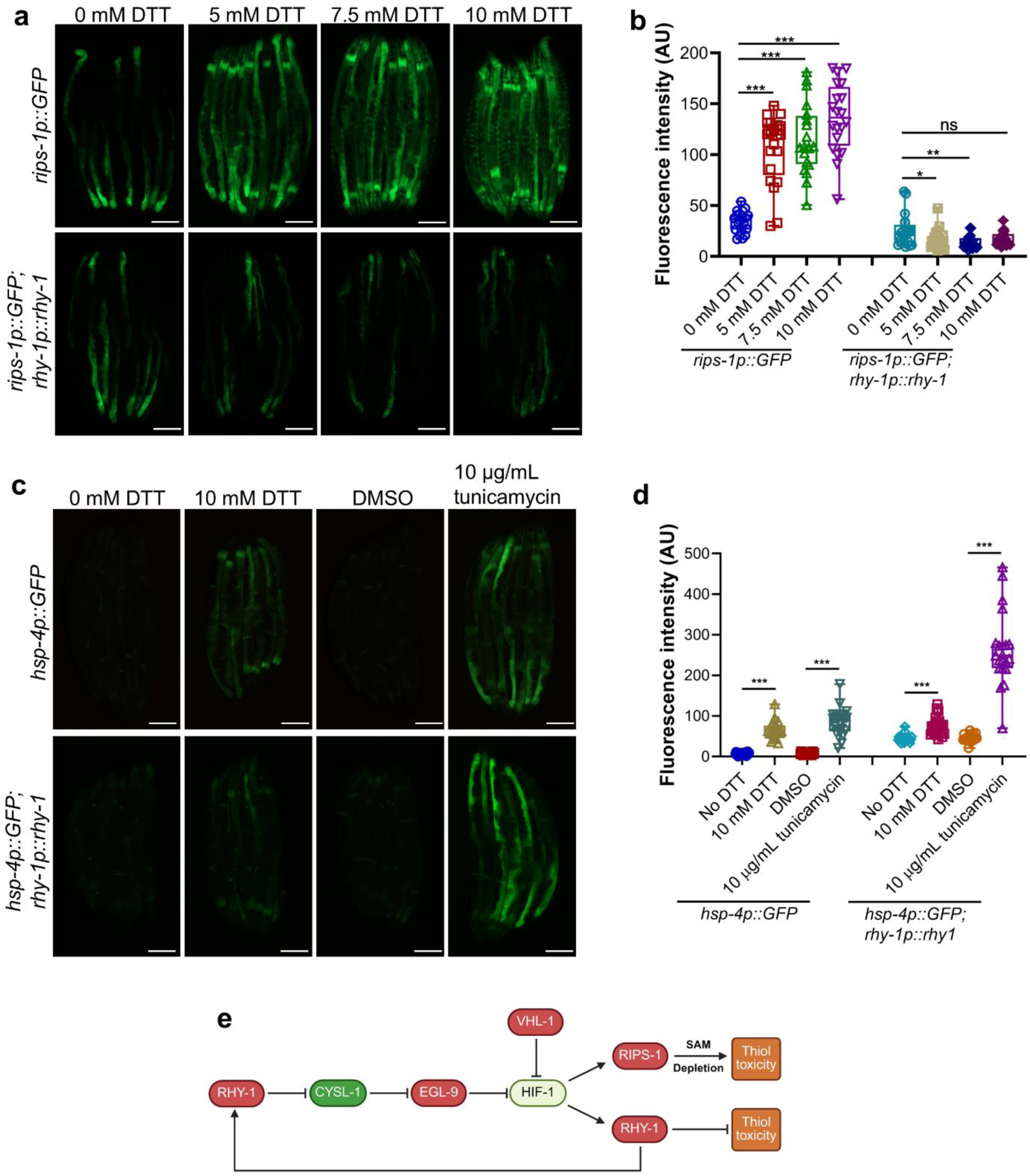
RHY-1 functions in an autoinhibitory feedback loop to protect from thiol toxicity. (a) Representative fluorescence images of *rips-1p::GFP* and *rips-1p::GFP;rhy-1p::rhy-1* worms after exposure of synchronized gravid adults to 0 mM, 5 mM, 7.5 mM, and 10 mM DTT for 12 hours. Scale bar = 200 μm. (b) Quantification of fluorescence levels of *rips-1p::GFP* and *rips-1p::GFP;rhy-1p::rhy-1* worms after exposure of synchronized gravid adults to 0 mM, 5 mM, 7.5 mM, and 10 mM DTT for 12 hours. \*\*\**p* < 0.001, \*\**p* < 0.01, \**p* < 0.05, and non-significant (ns) via ordinary one-way ANOVA followed by Dunnett’s multiple comparisons test (*n* = 20-21 worms each). (c) Representative fluorescence images of *hsp-4p::GFP* and *hsp-4p::GFP;rhy-1p::rhy-1* worms after exposure of synchronized gravid adults to 0 mM DTT, 10 mM DTT, DMSO control for tunicamycin, and 10 µg/mL tunicamycin for 20 hours. Scale bar = 200 μm. (d) Quantification of fluorescence levels of *hsp-4p::GFP* and *hsp-4p::GFP;rhy-1p::rhy-1* worms after exposure of synchronized gravid adults to 0 mM DTT, 10 mM DTT, DMSO control for tunicamycin, and 10 µg/mL tunicamycin for 20 hours. \*\*\**p* < 0.001 via the unpaired *t*-test (*n* = 21 worms each). (e) Model for the autoinhibitory feedback loop of the hypoxia response pathway that protects from thiol toxicity.

Overexpression of *rhy-1* could, in principle, suppress the hypoxia response pathway indirectly by promoting clearance of DTT and thereby preventing DTT-mediated activation of HIF-1. To test this possibility, we measured *hsp-4p::GFP* expression in worms overexpressing *rhy-1* following DTT exposure. HSP-4 is an ER-resident molecular chaperone whose expression is regulated by the ER unfolded protein response. DTT is known to induce *hsp-4* expression both by disrupting the methionine-homocysteine cycle and by causing protein misfolding in the ER (Gokul and Singh 2022). Overexpression of *rhy-1* increased basal *hsp-4p::GFP* levels but did not affect its induction by the ER stressor tunicamycin (Fig. 4c and d). Notably, the *hsp-4p::GFP* reporter exhibited partial induction upon DTT treatment in the *rhy-1* overexpression background (Fig. 4c and d), indicating that DTT continued to perturb cellular physiology despite elevated RHY-1 levels. These results suggested that *rhy-1* overexpression does not fully eliminate DTT and that suppression of the HIF-1 pathway reflects a direct inhibitory effect of RHY-1 rather than reduced DTT exposure. Together, these data indicated that RHY-1 protects against DTT toxicity and that increased RHY-1 levels prevent DTT-induced activation of the hypoxia response pathway. Overall, our findings support a model in which HIF-1 confers protection against thiol reductive stress through RHY-1, which in turn modulates both DTT toxicity and hypoxia signaling via an autoinhibitory feedback loop (Fig. 4e).

## Discussion

Our study identifies RHY-1 as a critical downstream effector of the hypoxia response pathway that protects *C. elegans* from DTT-induced reductive stress. Although RHY-1 was previously characterized as an ER membrane protein that negatively regulates HIF-1 under normoxic conditions (Shen et al. 2006), the physiological significance of its HIF-1-dependent upregulation remained unclear. Here, we demonstrate that RHY-1 is essential for survival under thiol reductive stress and fully rescues the DTT sensitivity of *hif-1* loss-of-function mutants. Thus, RHY-1 functions not only as a regulator of the hypoxia response pathway but also as an effector protein that mitigates thiol stress, an inducer of *rhy-1* expression itself through the hypoxia response pathway. This dual regulatory-effector role positions RHY-1 as a key component of an autoinhibitory feedback loop that modulates hypoxia signaling and thiol stress resistance (Fig. 4e).

RHY-1 has been implicated in modulating sensitivity to multiple thiol-containing molecules (Horsman et al. 2019; Warnhoff et al. 2024), although its precise function in thiol metabolism appears complex. In contrast to its role in DTT sensitivity, *rhy-1* loss-of-function mutants exhibit resistance to H_2_S and cysteine (Horsman et al. 2019; Warnhoff et al. 2024). These effects are mediated through HIF-1, suggesting that HIF-1 regulates distinct effector proteins that confer protection against specific thiol stressors. For example, exposure to H_2_S induces the HIF-1-dependent upregulation of the sulfide quinone oxidoreductase SQRD-1, which is required for defense against H_2_S toxicity (Budde and Roth 2011; Miller et al. 2011). Because protection from H_2_S and cysteine requires multiple downstream HIF-1 effectors, *rhy-1* loss-of-function mutants exhibit a protective phenotype by activating the hypoxia response pathway. In the case of DTT, however, RHY-1 overexpression fully restores the DTT sensitivity of *hif-1* mutants, indicating that RHY-1 is likely the principal HIF-1 effector responsible for protection from DTT toxicity. Interestingly, RHY-1 also protects from H_2_S toxicity in an HIF-1-independent but SKN-1-dependent manner (Horsman et al. 2019).

Increased activity of SKN-1, the *C. elegans* ortholog of mammalian NRF2, can upregulate *rhy-1* expression and compensate for *hif-1* loss under H_2_S exposure (Horsman et al. 2019). Furthermore, RHY-1 contributes to cysteine metabolism, as *rhy-1* mutants reduce sulfite production from cysteine and suppress toxicity in a sulfite-sensitized *suox-1(gk738847)* background (Warnhoff et al. 2024). Collectively, these findings suggest that RHY-1 plays multifaceted roles in both the regulation of hypoxia signaling and the metabolism of thiol-containing molecules. Notably, all three thiol-related stressors—H_2_S, cysteine, and DTT—are known activators of the hypoxia response pathway (Budde and Roth 2010; Ravi et al. 2023; Warnhoff et al. 2024).

A key mechanistic question emerging from our study is how RHY-1 confers protection from thiol toxicity. RHY-1 encodes a multipass ER membrane protein containing an acyltransferase-3 domain, which is typically associated with lipid biosynthesis and metabolic regulation (Shen et al. 2006). One possibility is that RHY-1 directly influences thiol metabolism. While S-acetylation of small-molecule thiols occurs in cells (Locigno et al. 2002; Filipovic et al. 2018; Di Paola et al. 2022), RHY-1 is unlikely to catalyze this reaction directly because it is predicted to possess O-acyltransferase rather than S-acetyltransferase activity. Nonetheless, *rhy-1* mutants display defects in cysteine catabolism and impaired sulfite production (Warnhoff et al. 2024), implying a direct metabolic role for RHY-1 in thiol processing. Alternatively, RHY-1 may indirectly mitigate thiol toxicity by altering cellular metabolism, for example through acylation of metabolic intermediates such as lipids or carbohydrates. The activation of the ER unfolded protein response in *rhy-1* overexpressing animals further suggests that RHY-1 is unlikely to function by directly clearing DTT, but instead may buffer the physiological disruptions caused by DTT. Future studies will be needed to elucidate the biochemical and metabolic mechanisms by which RHY-1 confers resistance to DTT.

In summary, RHY-1 emerges as a multifunctional regulator that integrates hypoxia signaling with thiol metabolism. By functioning in a negative feedback loop downstream of HIF-1, RHY-1 fine-tunes the hypoxia response while simultaneously providing protection from reductive stress. Future studies dissecting the biochemical activities of RHY-1 and its substrate specificity will be crucial for understanding how acyltransferases contribute to cellular adaptation under reductive stress conditions.

## Materials and methods

### Bacterial strains

*Escherichia coli* OP50 was used as the food source for *Caenorhabditis elegans* throughout this study. *E. coli* OP50 cultures were grown in Luria-Bertani broth at 37°C with shaking overnight.

### *C. elegans* strains and growth conditions

*C. elegans* hermaphrodites were maintained at 20°C on nematode growth medium (NGM) plates seeded with *E. coli* OP50 unless otherwise indicated. Bristol N2 was used as the wild-type control unless otherwise indicated. The following strains were used in the study: JSJ11 *rips-1(jsn11)*, JSJ13 *jsnIs1*[*rips-1p::GFP + myo-2p::mCherry*], JSJ14 *jsnIs1[rips-1p::GFP + myo-2p::mCherry];rhy-1(jsn13jsn14)*, JSJ15 *jsnIs1[rips-1p::GFP + myo-2p::mCherry];egl-9(jsn15)*, JSJ16 *jsnIs1[rips-1p::GFP + myo-2p::mCherry];vhl-1(jsn16)*, JSJ17 *jsnIs1[rips-1p::GFP + myo-2p::mCherry];cysl-1(jsn17)*, SJ4005 *zcIs4 [hsp-4::GFP]*, ZG31 *hif-1(ia4)*, JT307 *egl-9(sa307),* and RB1297 *rhy-1(ok1408)*. The following strains were generated by standard genetic crosses: *hif-1(ia4);rips-1(jsn11)*, *egl-9(sa307);rips-1(jsn11)*, *rhy-1(jsn13jsn14);rips-1(jsn11)*, *vhl-1(jsn16);rips-1(jsn11)*, and *rips-1(jsn11);cysl-1(jsn17)*.

### Generation of transgenic *C. elegans*

To overexpress *rhy-1*, the genomic region containing the *rhy-1* coding sequence and 995 bp of its upstream promoter was amplified from N2 genomic DNA. The fragment, including the stop codon, was cloned into the pPD95_77 plasmid using SalI and KpnI restriction sites. The primer sequences are listed in Table S1. The resulting construct (*rhy-1p::rhy-1*) was co-injected into N2 worms at 50 ng/µL, along with pCFJ104 (*myo-3p::mCherry*) as a co-injection marker (25 ng/µL). Transgenic worms carrying the extrachromosomal array *jsnEx5[rhy-1p::rhy-1 + myo-3p::mCherry]* were subsequently crossed into *hif-1(ia4)*, *rhy-1(jsn13jsn14)*, *rips-1p::GFP*, and *hsp-4p::GFP* backgrounds to generate *hif-1(ia4);jsnEx5[rhy-1p::rhy-1 + myo-3p::mCherry]*, *rhy-1(jsn13jsn14);jsnEx5[rhy-1p::rhy-1 + myo-3p::mCherry]*, *rips-1p::GFP;jsnEx5[rhy-1p::rhy-1 + myo-3p::mCherry]*, and *hsp-4p::*GFP*;jsnEx5[rhy-1p::rhy-1 + myo-3p::mCherry]*.

### Supplementation experiments

The following supplements were obtained from HiMedia BioSciences: DTT (#RM525) and vitamin B12 (cyanocobalamin) (#PCT0204). Tunicamycin (#654380) was purchased from Merck. Stock solutions were prepared in water, with DTT stocks freshly prepared prior to each experiment and vitamin B12 stocks stored at 4°C. Tunicamycin stock solutions were prepared in dimethyl sulfoxide (DMSO), and control plates contained an equivalent amount of DMSO. Stocks were diluted to the desired final concentrations in NGM prior to pouring the plates. NGM plates were seeded with *E. coli* OP50 and incubated at room temperature for at least three days before use.

### Development assays

To obtain synchronized *C. elegans* populations, 20-25 gravid adult hermaphrodites were transferred to NGM plates containing the indicated supplements and allowed to lay eggs for 2 hours. The adults were then removed, and the plates were incubated at 20°C for 72 hours. Worms were examined and scored at different developmental stages (L1/L2, L3, and L4/adult). Representative images were captured during assessment. Each condition was tested in at least three independent biological replicates.

### Fluorescence imaging and analysis

The fluorescence reporter strains were picked under a non-fluorescence stereomicroscope to minimize potential bias. Fluorescence imaging was carried out as described previously (Ghosh and Singh 2024; Das et al. 2025). Briefly, worms were anesthetized in 50 mM sodium azide in M9 buffer and mounted on 2% agarose pads. Fluorescence was visualized using a Nikon SMZ18 stereomicroscope, and images were analyzed using ImageJ software. Quantitative fluorescence intensity data were plotted using GraphPad Prism 8.

### Statistical analysis

Statistical analysis was performed using GraphPad Prism 8 as described earlier (Gahlot et al. 2025). Error bars represent mean ± standard deviation (SD). An unpaired, two-tailed, two-sample *t*-test was used when applicable, and the data were considered statistically significant when *p* < 0.05. For more than two samples, ordinary one-way ANOVA followed by Dunnett’s multiple comparisons test was used. In the figures, asterisks (*) denote statistical significance as follows: * for *p* < 0.05, **for *p* < 0.01, *** for *p* < 0.001, as compared with the appropriate controls. All experiments were performed in triplicate.

## Data availability

All data generated or analyzed during this study are included in the manuscript and supporting files.

## Acknowledgments

We thank the Caenorhabditis Genetics Center (funded by the NIH Office of Research Infrastructure Programs (P40 OD010440)) for providing the strains used in this study.

## Funding

This work was supported by the following grants: Ramalingaswami Re-entry Fellowship (Ref. No. BT/RLF/Re-entry/50/2020) and Har-Gobind Khorana-Innovative Young Biotechnologist Fellowship (File No. HRD-17011/2/2023-HRD-DBT) awarded by the Department of Biotechnology, India; Anusandhan National Research Foundation (ANRF) Core Research Grant (Ref. No. CRG/2023/001136) awarded by DST, India; STARS grant (File No. MoE-STARS/STARS-2/2023-0116) awarded by the Ministry of Education, India; Research Grant (Ref. No. 37/1741/23/EMR-II) awarded by the Council of Scientific & Industrial Research (CSIR), India; and IISER Mohali intramural funds.

## Declaration of interests

The authors declare no competing interests.

## Supplementary Figures

**Supplementary Figure 1.**
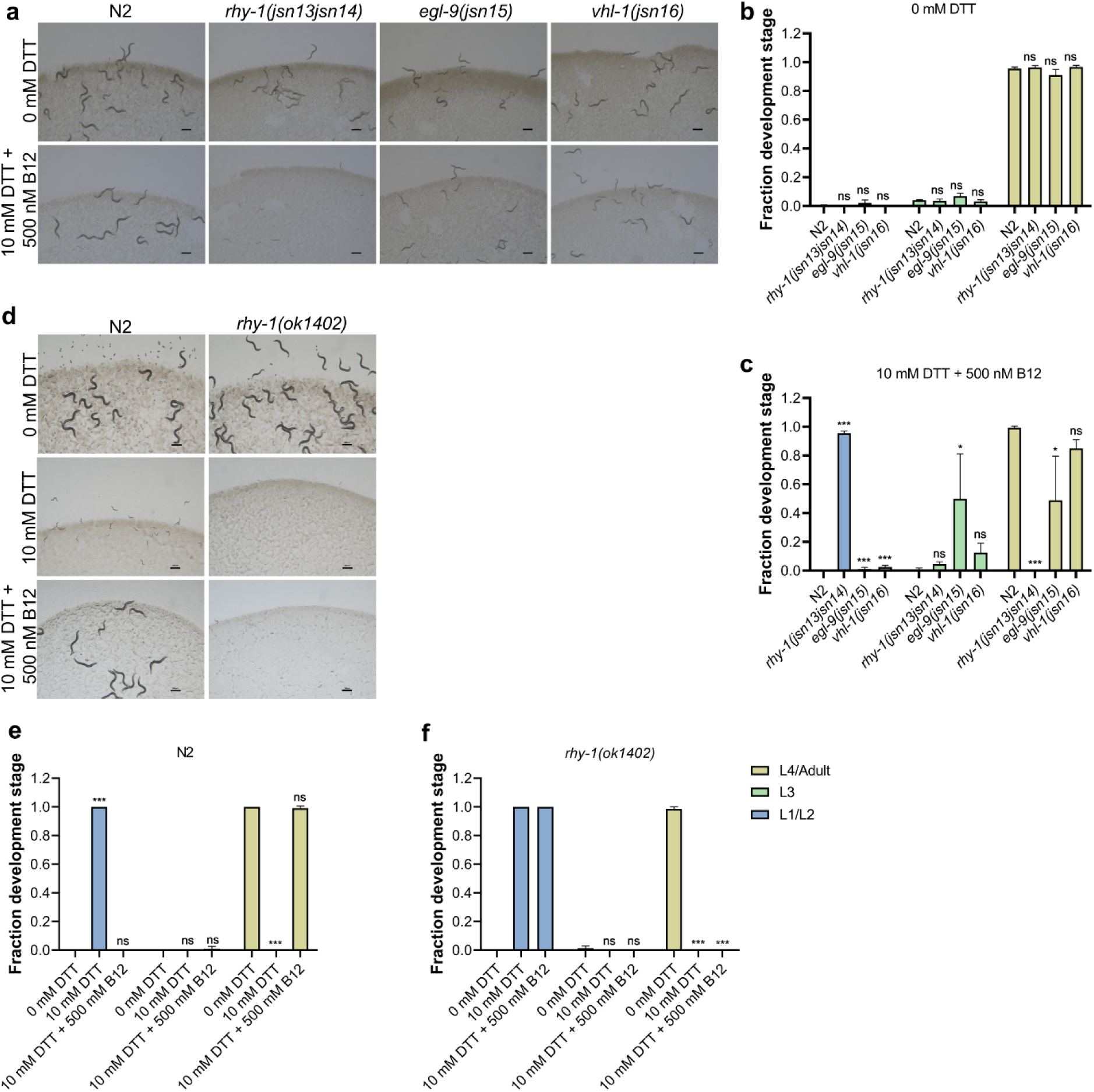
RHY-1 protects from thiol toxicity. (a) Representative images of N2, JSJ14 (*rhy-1(jsn13jsn14)*), JSJ15 (*egl-9(jsn15)*), and JSJ16 (*vhl-1(jsn16)*) worms exposed to 0 mM DTT or 10 mM DTT supplemented with 500 nM vitamin B12 on *E. coli* OP50 for 72 h at 20°C. Scale bar = 200 μm. (b)-(c) Quantification of different developmental stages of N2, JSJ14 (*rhy-1(jsn13jsn14)*), JSJ15 (*egl-9(jsn15)*), and JSJ16 (*vhl-1(jsn16)*) worms exposed to 0 mM DTT (b) or 10 mM DTT supplemented with 500 nM vitamin B12 (c) on *E. coli* OP50 for 72 h at 20°C. \*\*\**p* < 0.001, \**p* < 0.05, and non-significant (ns) via ordinary one-way ANOVA followed by Dunnett’s multiple comparisons test. For each developmental stage, the comparison is with N2 (*n* = 3 biological replicates; worms per condition per replicate > 80). (d) Representative images of N2 and *rhy-1(ok1402)* worms exposed to 0 mM DTT, 10 mM DTT, or 10 mM DTT supplemented with 500 nM vitamin B12 on *E. coli* OP50 for 72 h at 20°C. Scale bar = 200 μm. (e)-(f) Quantification of different developmental stages of N2 (e) and *rhy-1(ok1402)* (f) worms exposed to 0 mM DTT, 10 mM DTT, or 10 mM DTT supplemented with 500 nM vitamin B12 on *E. coli* OP50 for 72 h at 20°C. \*\*\**p* < 0.001 and non-significant (ns) via ordinary one-way ANOVA followed by Dunnett’s multiple comparisons test. For each developmental stage, the comparison is with the 0 mM DTT control (*n* = 3 biological replicates; worms per condition per replicate > 55). For panels b, c, e, and f, the different developmental stage symbols are shown in panel f.

**Supplementary Figure 2.**
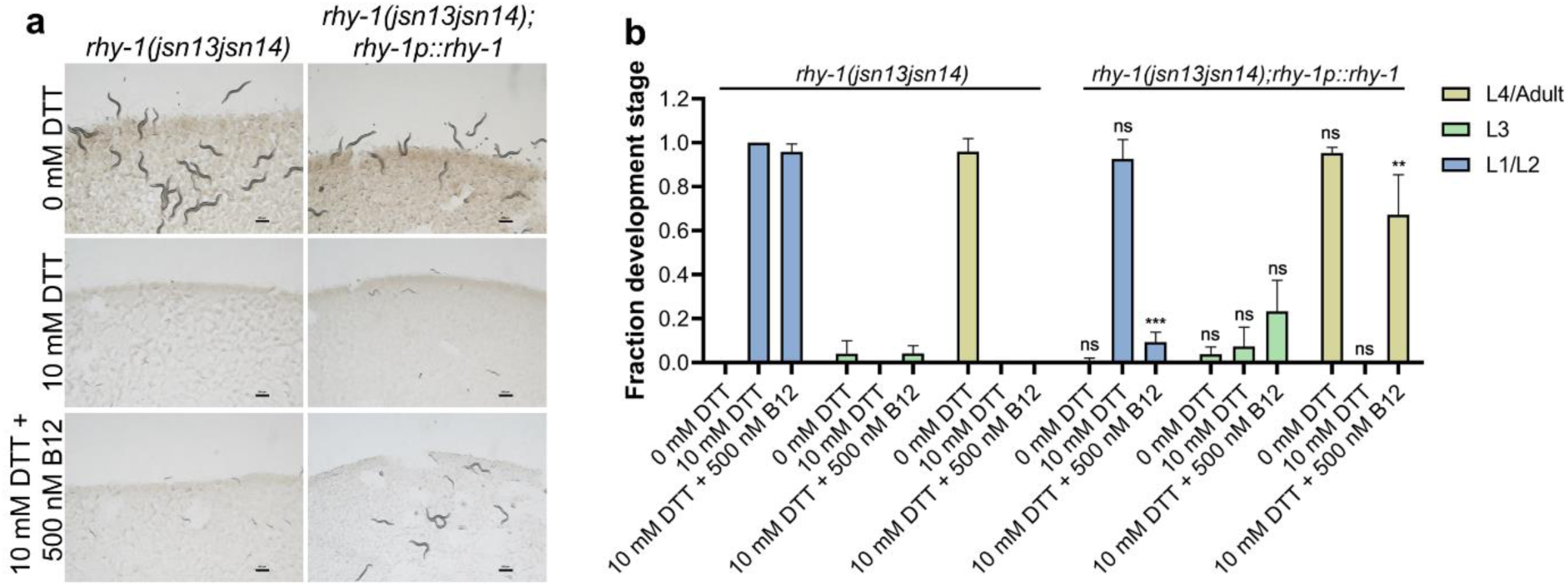
RHY-1 protects from thiol toxicity. (a) Representative images of *rhy-1(jsn13jsn14)* and *rhy-1(jsn13jsn14);rhy-1p::rhy-1* worms exposed to 0 mM DTT, 10 mM DTT, or 10 mM DTT supplemented with 500 nM vitamin B12 on *E. coli* OP50 for 72 h at 20°C. Scale bar = 200 μm. (b) Quantification of different developmental stages of *rhy-1(jsn13jsn14)* and *rhy-1(jsn13jsn14);rhy-1p::rhy-1* worms exposed to 0 mM DTT, 10 mM DTT, or 10 mM DTT supplemented with 500 nM vitamin B12 on *E. coli* OP50 for 72 h at 20°C. \*\*\**p* < 0.001, \*\**p* < 0.01, and non-significant (ns) via unpaired t-test. For each condition of *rhy-1(jsn13jsn14);rhy-1p::rhy-1* worms, the comparison is with the corresponding *rhy-1(jsn13jsn14)* condition (*n* = 3 biological replicates; worms per condition per replicate > 35).

**Table S1:**
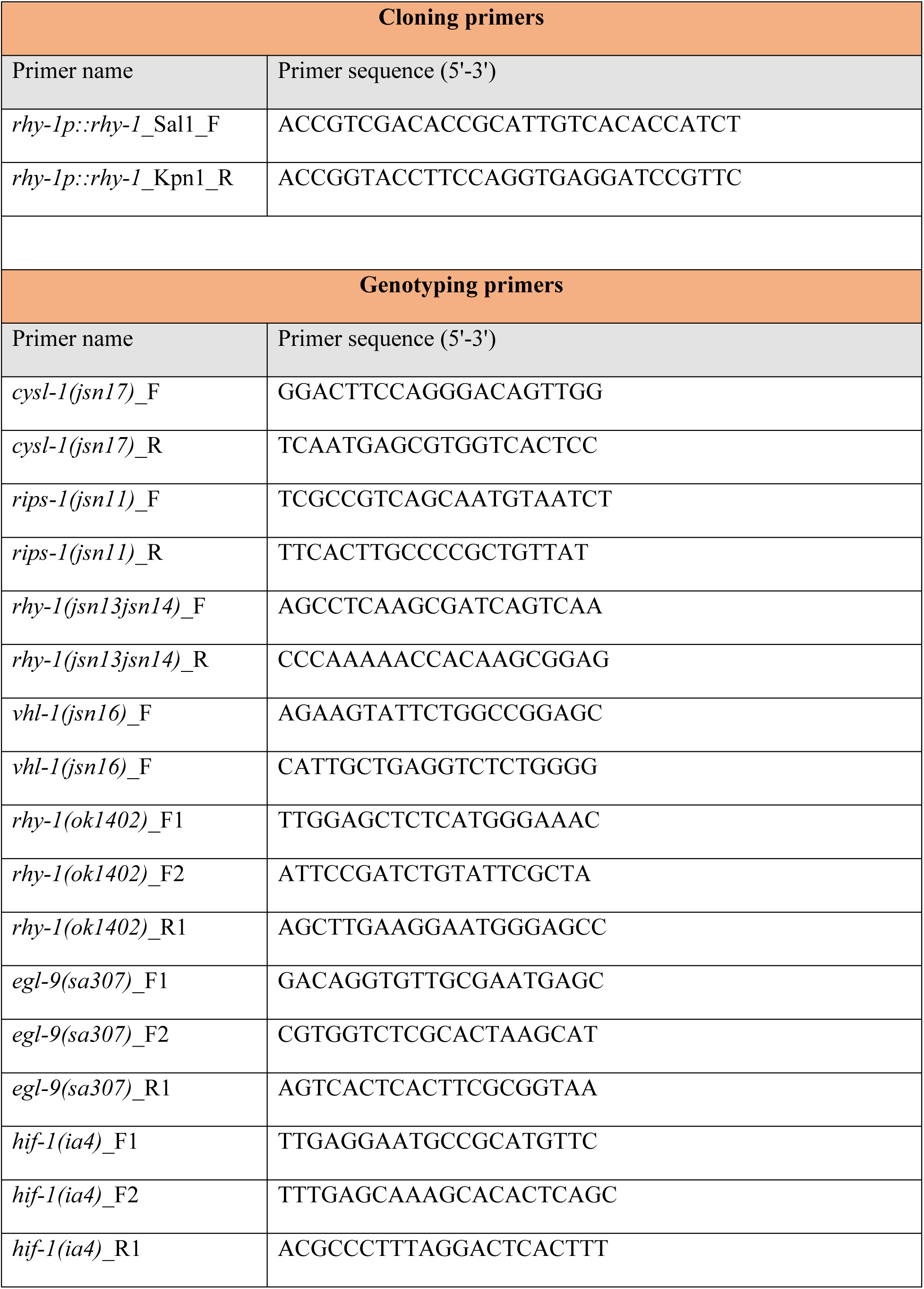
Primers used in the study.

## Notes

### Competing Interest Statement

The authors have declared no competing interest.

### Summary of Updates

The text has been revised. The bimolecular fluorescence complementation (BiFC) assay experiments have been removed. Data with hsp-4p::GFP strain are added under rhy-1p::rhy-1 overexpression conditions.

